# Independent component analysis reveals the transcriptional regulatory modules in *Bradyrhizobium diazoefficiens* USDA110

**DOI:** 10.1101/2023.06.30.547077

**Authors:** Zhi-Peng Gao, Wei-Cheng Gu, Jie Li, Qin-Tian Qiu, Bin-Guang Ma

**Author notes:** Corresponding author., E-mail address (Bin-Guang Ma).

## Abstract

The dynamic adaptation of bacteria to environmental changes is achieved through the coordinated expression of many genes, which constitutes a transcriptional regulatory network (TRN). *Bradyrhizobium diazoefficiens* USDA110 is an important model strain for the study of symbiotic nitrogen fixation (SNF), and its SNF ability largely depends on the TRN. In this study, independent component analysis was applied to 226 high-quality gene expression profiles of *B. diazoefficiens* USDA110 microarray datasets, from which 64 iModulons were identified. Using these iModulons and their condition-specific activity levels, we (1) provided new insights into the connection between the FixLJ-FixK_2_-FixK_1_ regulatory cascade and quorum sensing, (2) discovered the independence of the FixLJ-FixK_2_-FixK_1_ and NifA/RpoN regulatory cascades in response to oxygen, (3) identified the FixLJ-FixK_2_ cascade as a mediator connecting the FixK_2_-2 iModulon and the Phenylalanine iModulon, (4) described the differential activation of iModulons in *B. diazoefficiens* USDA110 under different environmental conditions, and (5) proposed a notion of active-TRN based on the changes in iModulon activity to better illustrate the relationship between gene regulation and environmental condition. In sum, this research offered an iModulon-based TRN for *B. diazoefficiens* USDA110, which formed a foundation for comprehensively understanding the intricate transcriptional regulation during SNF.

## 1. Introduction

*Bradyrhizobium diazoefficiens* USDA110 (formerly known as *Bradyrhizobium japonicum* USDA110) [1] is an important model strain for the study of symbiosis. It can establish symbiotic relationship with *Glycine max*, *Vigna unguiculata*, *Vigna radiata*, *Macroptilium atropurpureum* [2] and has strong ability of symbiotic nitrogen fixation (SNF). It is widely used in agriculture and ecological fields [3]. *B. diazoefficiens* USDA110 forms a symbiotic system with legumens, mainly involving the following processes: 1) *B. diazoefficiens* USDA110 receives the signal molecules from plant roots and releases nodulation factors; 2) *B. diazoefficiens* USDA110 enters the soybean roots and differentiates into bacteroids, which are then wrapped by functional symbiotic membranes to form a low-oxygen environment; 3) *B. diazoefficiens* USDA110 expresses the related genes of nitrogenase complex and initiates the nitrogen fixation process [4]. Moreover, *B. diazoefficiens* USDA110 also has denitrification ability [5]. The correct implementation of this series of complex biological processes depends on the orderly expression among genes. A comprehensive understanding of the transcriptional regulatory network (TRN) of *B. diazoefficiens* USDA110 can help us uncover the molecular mechanisms related to SNF.

In recent years, there has been great progress in the omics research of *B. diazoefficiens* USDA110 [6–8]. The number of databases related to rhizobia omics is increasing, and the relevant gene annotation information and multi-omics data are gradually complete. Based on these data, Yang et al. established the genome-scale metabolic network of the *B. diazoefficiens* USDA110 [9] and Ma et al. constructed its protein-protein interaction network [10]. However, there are rare studies on the TRN of *B. diazoefficiens* USDA110, and related databases such as RhizoRegNet [11], PRODORIC [12], RegPrecise [13], RegTransBase [14], and RhizoBindingSites [15] have incomplete information and lack relevant connections between gene regulation and environmental conditions.

Independent component analysis (ICA) [16] is an unsupervised machine learning algorithm, which is particularly suitable for blind source separation of mixed signals. ICA has been used in bacterial transcriptomic datasets to identify independent sets of regulated genes and their regulatory factors [17–20]. Briefly, by decomposing the gene expression matrix through ICA, we can obtain groups of genes (called iModulons) that represent independent regulatory signals. These genes may be regulated by the same or related regulators [20]. Each iModulon has a corresponding activity in different conditions, which reflects the number of iModulon genes present under that condition. Unlike regulons [21], which are obtained from experimental data by analyzing transcription factor (TF) binding sites (TFBS) in a bottom-up fashion, iModulons are calculated based on gene expression via a top-down way [22].

Researchers have generated a lot of meaningful hypotheses by applying the iModulon analysis process to different species. Chauhan et al. [23] applied the iModulon analysis process to *Sulfolobus acidocaldarius* and predicted a DNA export system composed of five uncharacterized genes. Lim et al. [17] compiled the gene expression profiles of *Pseudomonas putida* KT2440 during growth and stationary phases, used iModulons to describe the transition of transcriptional levels from the former to the latter, and revealed multiple evolutionary strategies for achieving rapid growth rate in this species. Rajput et al. [18] compiled the gene expression profiles of *Pseudomonas aeruginosa* under different nutrient conditions and antibiotic treatments, and then used ICA to analyze the iModulon structure, resulting in new roles of transcriptional regulators in antibiotic efflux regulation and the association between nutrient substrates and expression of antibiotic efflux pump genes. By now, the iModulon structures of 11 species have been analyzed, and related datasets and results are present in the iModulonDB database [24].

Currently, most studies on the transcriptional regulatory relationships of *B. diazoefficiens* USDA110 were mainly carried out by analyzing transcriptome differences in gene-deficient strains, combined with motif search for TFBS in promoter region [25–29]. Socorro Mesa et al. [30] studied the FixLJ-FixK_2_-FixK_1_ cascade of *B. diazoefficiens* USDA110 and found that FixJ positively regulates FixK_2_ at 0.5% oxygen concentration. Analysis of the differences in bacterial transcription levels by microarray showed that 204 genes were affected by FixK_2_ in *B. diazoefficiens* USDA110, and this number further increased 21 days after bacteroid formation; some of the affected genes are even not regulated by FixJ. These results indicate that the transcriptional regulation of *B. diazoefficiens* USDA110 is very complex, and the target genes regulated by the same transcription factor may vary with environmental changes. Although some transcriptional regulatory relationships can be accurately obtained using experimental methods, they cannot provide a full picture for the transcriptional regulation of *B. diazoefficiens* USDA110 under different environmental conditions.

To better understand the TRN of *B. diazoefficiens* USDA110, we compiled a microarray dataset based on the GPL3401 microarray platform, which contains 226 *B. diazoefficiens* USDA110 transcriptome samples. Through ICA algorithm, 64 iModulons were extracted and they were used to extend the existing regulons. We provided new insights into the relationship between the FixLJ-FixK_2_-FixK_1_ regulatory cascade and quorum sensing. We described the independence of the FixLJ-FixK_2_-FixK_1_ and NifA/RpoN cascades in response to oxygen environment. We identified the differential activation states of iModulons under different environments in *B. diazoefficiens* USDA110, and based on this, proposed the notion of active-TRN, demonstrating the relevant connection between gene regulation and environmental condition.

## 2. Materials and methods

### 2.1 Data collection and preprocessing

We downloaded all the microarray data of Affymetrix custom *Bradyrhizobium japonicum* strain USDA110 19K array (Platform GPL3401) from the GEO database, which covers 23 series. Samples with poor replicability (sample correlation coefficient < 0.95) were removed, and the final dataset contained 226 samples involving 54 unique experimental conditions. The median Pearson correlation coefficient between sample replicates is 0.96 (Figure S1A). We performed log2 transformation on the data. In order to minimize the noise caused by batch effects among samples, only the samples from two institutions (ETH Zurich and TU Dresden) were used in the dataset (Figure S1B). The data have been verified by Principal Components Analysis (PCA) and have high self-consistency between the samples.

At the same time, we also selected a reference condition for each project, which was usually the control group in the experiment design. Following a previous study [31], we normalized the data by subtracting the mean value of the corresponding reference condition from all expression data in the project. Anand V. Sastry et al.released the procedure for analyzing iModulon (https://github.com/avsastry/modulome-workflow), and we referred to their public code and made some modifications for our research (https://github.com/mbglab/BdTRNiMod).

### 2.2 Independent component analysis

Following the PyModulon workflow, we used the well-organized dataset as input and the scripts provided by Anand V. Sastry et al. (with modifications) to decompose the expression matrix into two matrices, A and M [32]. Briefly, the scikit-learn (v0.23.2) [33] implementation of FastICA [34] was performed 100 times with a convergence tolerance of 10^-7^ to obtain robust independent components (ICs). The resulting independent components (ICs) were clustered using DBSCAN [35] to identify robust ICs, with an epsilon of 0.1 and minimum cluster seed size of 50. To account for identical components with opposite signs, the following distance metric was used for computing the distance matrix: *d_x_*, *_y_* = 1 - || *ρ_x_*, *_y_* ||, where *ρ_x_*, *_y_* is the Pearson correlation coefficient between components *x* and *y*. The final robust ICs were defined as the centroids of the cluster. We performed clustering of the gene expression profile multiple times for dimensions between 20 and 215 with a step size of 5. The optimal dimension of 210 was finally chosen (Figure S2).

### 2.3 Characterization of iModulons

We obtained 64 iModulons and determined the iModulon gene sets using functions provided by PyModulon. The RefSeq ID of *B. diazoefficiens* USDA110 genome was BA000040.2, and the gene annotation came from the NCBI Prokaryotic Genome Annotation Pipeline. In addition, KEGG [36] and Cluster of Orthologous Groups (COG) [37] information was obtained using EggNOG mapper [38]. Operon information was obtained from BioCyc [39]. The gene ontology (GO) annotation came from AmiGO2 [40]. The draft TRN of *B. diazoefficiens* USDA110 was constructed based on the RegPrecise database [13] and our manually collected data from the literature. To determine the overlap between iModulons and regulons, we performed TF enrichment analysis, and the relevant method was described as in Yoo et al. [31]. We used "iModulon recall" and "regulon recall" values to evaluate the degree of overlap between predicted iModulons and regulons. "iModulon recall" represents the proportion of shared genes and genes in an iModulon, while "regulon recall" is the proportion of shared genes and genes in a regulon [20]. Common motifs in the upstreams of iModulons were enriched by using the find_motifs function that uses MEME [41] with a maximum window of 30 bp. When motifs were compared, the web-based TOMTOM [42] was utilized. As the statistical significance, *E*-values were provided as adjusted *p*-values when more than one common sequence was identified or compared.

### 2.4 Active-TRN construction

We constructed active-TRN based on the differences in iModulon activity. Firstly, we extracted the activity for each iModulon from matrix A under the aerobic condition of free-living *B. diazoefficiens* USDA110 as the reference state, and then identified the activity difference between the reference state and the target environmental condition by comparing the distributions of corresponding iModulon activities with a two-sample Kolmogorov-Smirnov test. Then, we selected the iModulons that showed a significant increase in activity under the target environmental condition. Finally, we extracted the regulatory relationships involved in these iModulons to construct the active-TRN.

## 3. Results

### 3.1 Independent component analysis revealed the independent signal modules of the *B. diazoefficiens* USDA110 transcriptome

We used the ICA algorithm to decompose the gene expression matrix and obtained 64 independent components (iModulons). These iModulons contain a total of 2269 genes, accounting for 27.6% of the annotated genes, and explain 81.1% of the gene expression variation. To identify the features of iModulons, we first constructed the draft TRN of *B. diazoefficiens* USDA110 based on published literature and the RegPrecise database [13]. The draft TRN includes 633 regulatory relationships involving 43 regulators. Next, we compared each iModulon with known regulons and found that 13 iModulons significantly overlapped with existing regulons. We examined the degree of overlap between iModulons and regulons by calculating regulon recall (RR) and iModulon recall (MR); the degree of overlap was divided into four categories with a threshold value of 0.6 for both RR and MR: well-matched, regulon subset, regulon discovery, and poorly-matched [17]. The results are shown in Figure 1A, where the HutC iModulon shows a high degree of overlap with the known HutC regulon, indicating that it accurately captures the previously reported regulation of 9 target genes by the histidine utilization repressor HutC (blr6245). Among these target genes, blr6249 and blr6250 are not in the currently known HutC regulon (Figure 1B). Through annotation information, it was found that these two genes (blr6249, blr6250) and the two genes (blr6247, blr6248) in the HutC regulon belong to the same transcriptional unit. It can be assumed that all the above four genes are regulated by HutC. Therefore, the HutC regulon has been successfully expanded by the HutC iModulon. After analyzing the upstream promoter regions of the HutC iModulon using MEME [41], three motifs were obtained (Figure S3), among which motif-2 (Figure S3B) and motif-3 (Figure S3C) were enriched in all the four promoter regions, suggesting that these motifs are potential binding sites for the HutC TF. In addition, the activity of the HutC iModulon decreased under anaerobic conditions (Figure 1C), which is consistent with previous studies [43]. Therefore, the iModulon effectively reflects gene regulatory relationships and can supplement the existing TRN.

**Figure 1.**
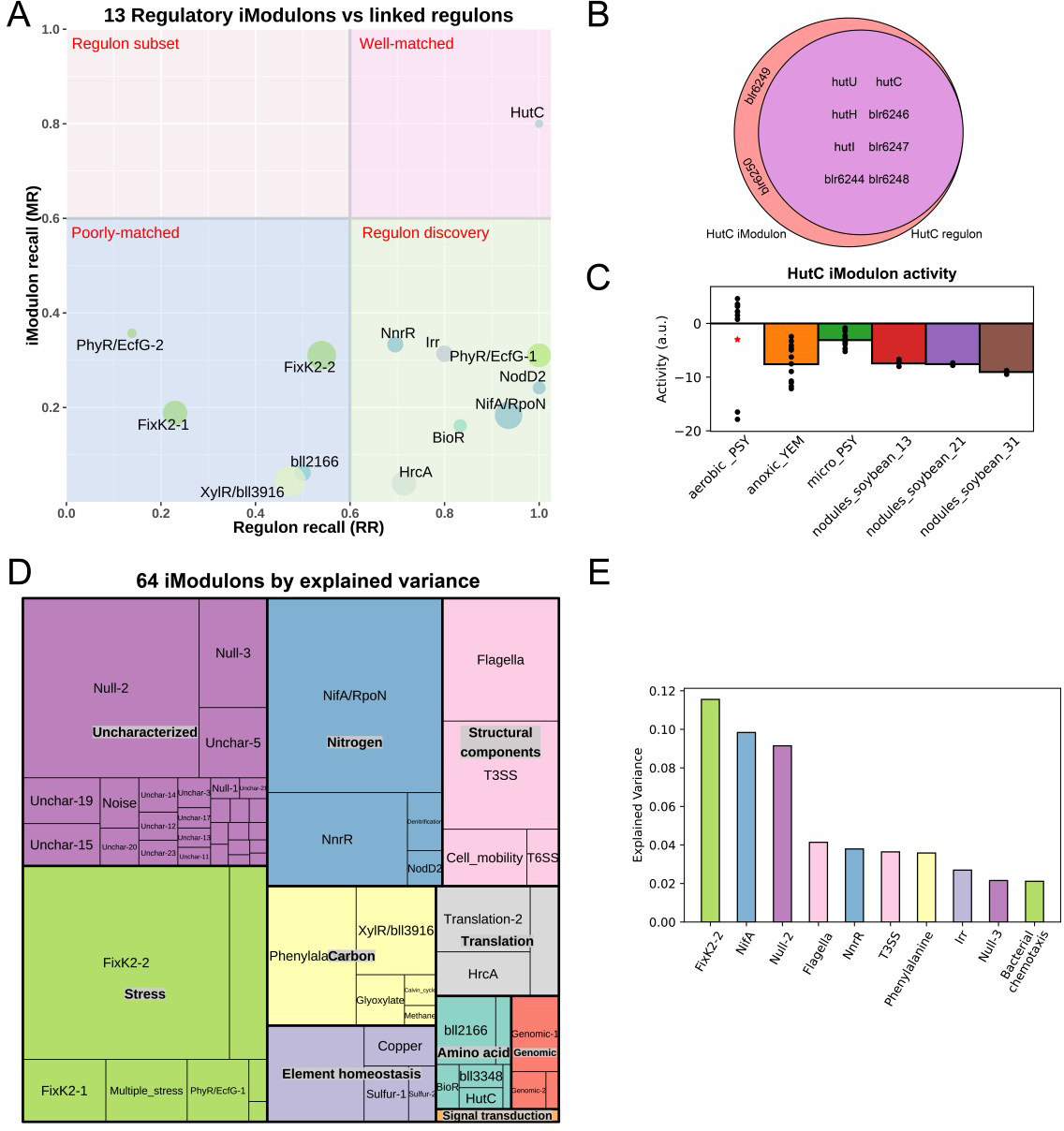
The compendium of Independent Component Analysis (ICA) for the *B. diazoefficiens* USDA110 transcriptome. (A) The scatter plot shows Regulon Recalls (RR) and iModulon Recalls (MR) for the 13 regulatory iModulons. (B) Comparison of genes in the reported HutC regulon and the HutC iModulon. (C) HutC iModulon activities under anoxic conditions. Asterisks indicate the reference condition (aerobic with PSY media). (D) A treemap of *B. diazoefficiens* USDA110’s 64 iModulons. The size of each rectangle indicates the explained variances in the data sets attributable to each iModulon. (E) Ten iModulons with the highest explained variances.

The remaining 51 iModulons were characterized by information from public databases and literature. Adopting the scripts provided by Lim et al. [17], we classified these iModulons into 10 functional categories based on the biological processes and functions they were involved in (Figure 1D). The FixK_2_-2 iModulon, which is related to the oxygen adaptation mechanism, made the largest contribution to gene expression variances, indicating that the sense of oxygen environment is the most important factor affecting the biological functions of *B. diazoefficiens* USDA110. Other iModulons with significant contributions to the expression variances were mainly involved in SNF (NifA/RpoN iModulon and NnrR iModulon), cell mobility (Flagella iModulon), and secretion system (T3SS iModulon) (Figure 1E), which reflects the main biological characteristics of *B. diazoefficiens* USDA110 and its physiological states under specific environmental conditions.

### 3.2 The FixK_2_ iModulon illustrates the regulatory cascade of FixK_2_ and implies its relationship with quorum sensing

As the FixK_2_-2 iModulon had the greatest contribution to gene expression variances, we performed extensive analysis to identify and characterize it. FixK_2_ is a CRP/FNR-type TF that plays a central role in the adaptation of *B. diazoefficiens* USDA110 to low-oxygen environments, SNF, and denitrification processes [28, 44]. FixK_2_ is regulated by a two-component system FixLJ, in which FixL senses the oxygen concentration in the environment and phosphorylates FixJ under a condition of less than 5% oxygen concentration, and the phosphorylated FixJ activates FixK_2_. Then, FixK_2_ activates the microaerobic adaptive operon *fixNOQP* and *fixGHIS*, several heme synthesis genes (*hemA*, *hemB*, *hemN_1_*, *hemN_2_*), denitrification genes (*nnrR*, *napEDABC*, *nirK*, *norCBQD*, *nosRZDYFLX*), some hydrogen oxidation genes (*hup* genes) [30], and the *fixK_1_* gene that is homologous to *fixK_2_*. The FixK_2_-2 iModulon includes 174 genes, most of which are related to low-oxygen adaptation, and 70 genes overlap with the FixK_2_ regulon, accounting for 58.3% of the FixK_2_ regulon genes (Figure 2A). The activity of FixK_2_-2 iModulon significantly increases in microaerobic environments and symbiotic nodulation state (Figure 2B), indicating that the FixK_2_-2 iModulon of *B. diazoefficiens* USDA110 plays a key role in the adaptation of low-oxygen environment, which is consistent with the main function of FixK_2_ regulon.

**Figure 2.**
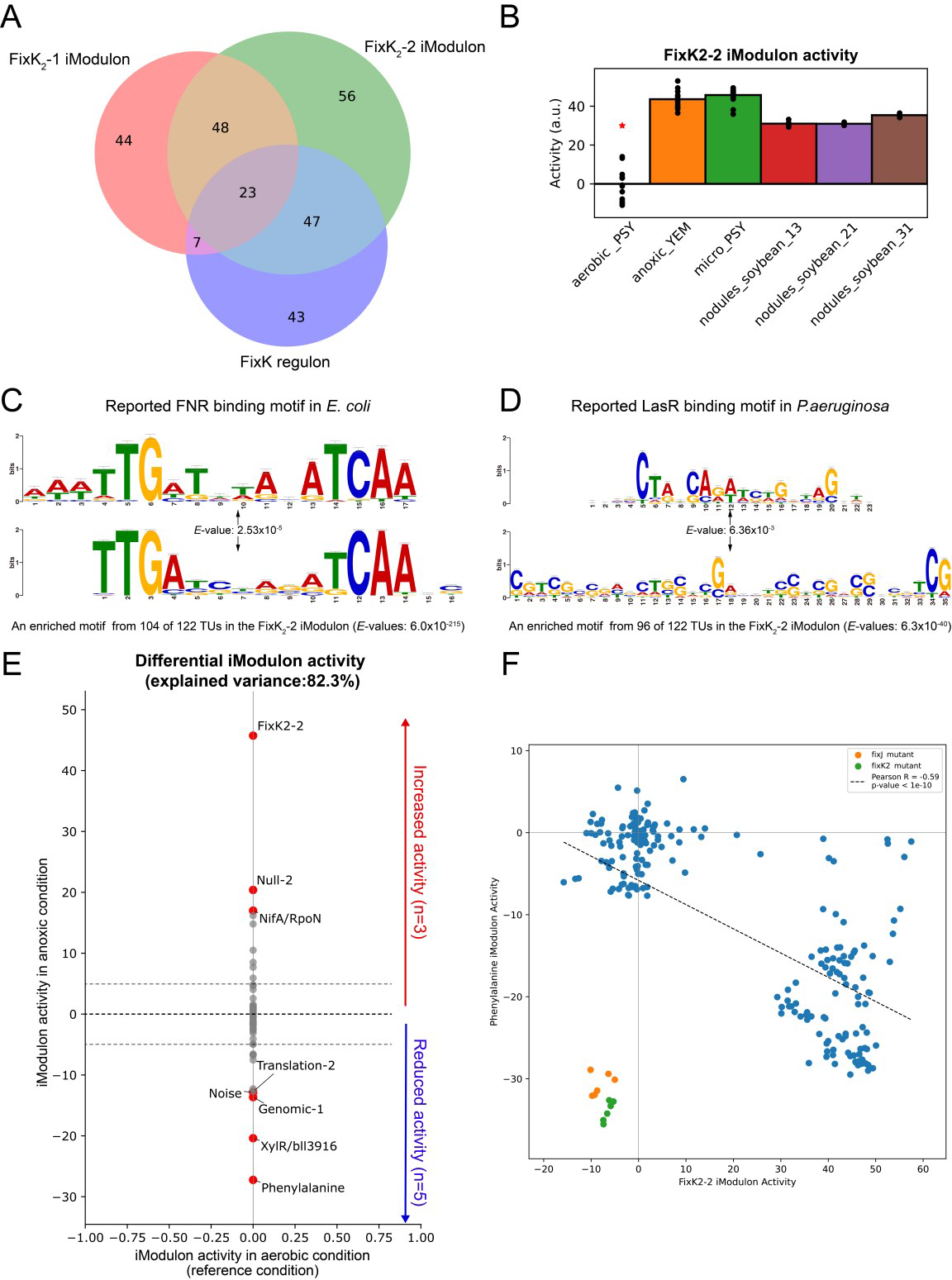
Overview of the FixK_2_ iModulon. (A) Comparison of genes in the reported FixK regulon, FixK_2_-1 iModulon and FixK_2_-2 iModulon. (B) FixK_2_-2 iModulon activities under anoxic condition. (C) Comparison of a motif found in the upstream regions of the FixK_2_-2 iModulon genes and the previously known binding motif of FNR in *Escherichia coli*. (D) Comparison of a motif found in the upstream regions of the FixK_2_-2 iModulon genes and the previously known binding motif of LasR in *P. aeruginosa*. (E) Differentially changed iModulon activities between anoxic condition and aerobic condition (threshold = 5 and *FDR* = 0.05). (F) Correlation of activities between the FixK_2_-2 iModulon and the Phenylalanine iModulon.

Through MEME enrichment analysis for the upstream sequence of all transcriptional units of the FixK_2_-1 and FixK_2_-2 iModulons member genes, we identified a motif that is highly consistent with the known FixK_2_ binding site (Figure 2C). Since FixK_2_ belongs to the Fnr family, this binding site is highly similar to the Fnr binding site of *Escherichia coli*. The FixK_2_-1 iModulon involves 84 operons, 47 of which contain this motif; the FixK_2_-2 iModulon involves 122 operons, 104 of which contain this motif. By comparing with the FixK_2_ regulon, we found 148 genes that do not belong to the FixK_2_ regulon (Figure 2A), and these genes may be potential targets of FixK_2_, which requires further validation. It is worth noting that the activities of the FixK_2_-1 and FixK_2_-2 iModulons significantly decreased in *fixJ* and *fixK_2_* mutant strains (Figure S4), indicating that *fixJ* and *fixK_2_* are the key TFs regulating the gene expression in FixK_2_-1 and FixK_2_-2 iModulons.

By comparing the activities of the FixK_2_-1 and FixK_2_-2 iModulons, we found that the activity of the FixK_2_-2 iModulon was significantly higher than that of the FixK_2_-1 iModulon in the blr1880 project. After the deletion of the *blr1880* gene, the FixK_2_-1 iModulon activity was significantly decreased (Figure S5). Blr1880 is a LuxR family TF and this family of proteins is usually associated with quorum sensing [45]. We hypothesized that the difference in activity between the FixK_2_-1 and FixK_2_-2 iModulons is due to the quorum sensing function mediated by blr1880. To verify this hypothesis, we annotated the enriched motifs using TOMTOM and found that a common motif “CGTCGCSGHSCTSSWCGABVKSSTCGMSSHSVTCG” in the FixK_2_-1 and FixK_2_-2 iModulons is similar to the LasR TFBS motif in *Pseudomonas aeruginosa* (*p*-values of 1.08×10^-5^ and 1.75×10^-5^, respectively) (Figure 2D). The LasR TF belongs to the LuxR protein family and regulates quorum sensing in *Pseudomonas aeruginosa* [46]. This suggests that these sites are potential binding sites for blr1880, indicating that FixK_2_ and blr1880 have interrelated regulatory functions. Given that FixK_2_ is regulated by oxygen and blr1880 primarily participates in quorum sensing, it is worthwhile to investigate whether the oxygen concentration also impacts quorum sensing in *B. diazoefficiens* USDA110. By examining the gene composition of the FixK_2_-1 and FixK_2_-2 iModulons, we found that the FixK_2_-2 iModulon contains more genes related to denitrification, including *napABCDE* and *nnrR*, while the FixK_2_-1 iModulon does not. This indicated that the FixK_2_-2 iModulon better represents the FixLJ/FixK_2_/NnnR cascade, while the FixK_2_-1 iModulon better represents the blr1880-involved quorum sensing. Overall, these two iModulons clearly demonstrate that FixK_2_ is involved in anaerobic adaptation and denitrification processes in *B. diazoefficiens* USDA110, and imply a potential link between the genes involved in microaerobic adaptation and those involved in quorum sensing. This provides a new clue for further understanding the relationship between these biological processes.

Next, by comparing the activities of iModulons under aerobic and anaerobic conditions, 8 iModulons with significant differences were obtained, among which 3 iModulons were significantly up-regulated and 5 iModulons were significantly down-regulated. The largest activity changes are shown in FixK_2_-2 iModulon and Phenylalanine iModulon (Figure 2E). The Phenylalanine iModulon contains 25 genes, and the genes with high weights mainly include hydroxypropanoate isomerase (*hyi*), aldehyde dehydrogenase (bll4784), and cytochrome c-type (blr7488), which play crucial roles in carbohydrate metabolism and energy production. Therefore, the decreased activity of Phenylalanine iModulon indicates that carbohydrate metabolism is suppressed, resulting in reduced energy production and inhibition of basic cellular metabolism under anaerobic conditions. In most cases, the activities of FixK_2_-2 iModulon and Phenylalanine iModulon are negatively correlated significantly (Figure 2F), while this negative correlation disappeared in the *fixJ* mutant and the *fixK_2_* mutant, indicating that FixLJ/FixK_2_ may act as a mediator connecting the FixK_2_-2 and Phenylalanine iModulons.

### 3.3 The NifA/RpoN iModulon successfully captures nitrogen-fixing gene cluster

NifA is one of the most important TFs in *B. diazoefficiens* USDA110, and its target genes include the *nif* gene cluster directly involved in nitrogenase synthesis, as well as the *fixRnifA* operon, which is subject to NifA-dependent autoregulation under low-oxygen conditions. It also regulates genes involved in SNF such as *fixA*, *fixBCX*, *nif*, *fix*, *hup* operons, *groESL*, *fdxN*, and *rpoN*, as well as genes with unknown functions such as *nrgA* and *nrgBC* [47, 48]. Studies have shown that NifA is also a necessary gene for the maximal expression of denitrification genes in *B. diazoefficiens* USDA110 [5]. NifA cooperates with RNA polymerase containing sigma 54 to activate gene expression. In *B. diazoefficiens* USDA110, sigma 54 is encoded by two highly similar and functionally equivalent genes (*rpoN_1_* and *rpoN_2_*) [49]. The NifA/RpoN iModulon includes 158 genes, most of which are located in a region approximately 2 Mb from the start site of DNA replication, and this region mainly contains genes directly related to SNF (Figure 3A). By enriching the upstream promoter regions of the NifA/RpoN iModulon using MEME, a motif highly similar to the *Vibrio cholerae* RpoN TFBS motif was found (Figure 3B). Therefore, the above results are consistent with the functional characteristics of NifA.

**Figure 3.**
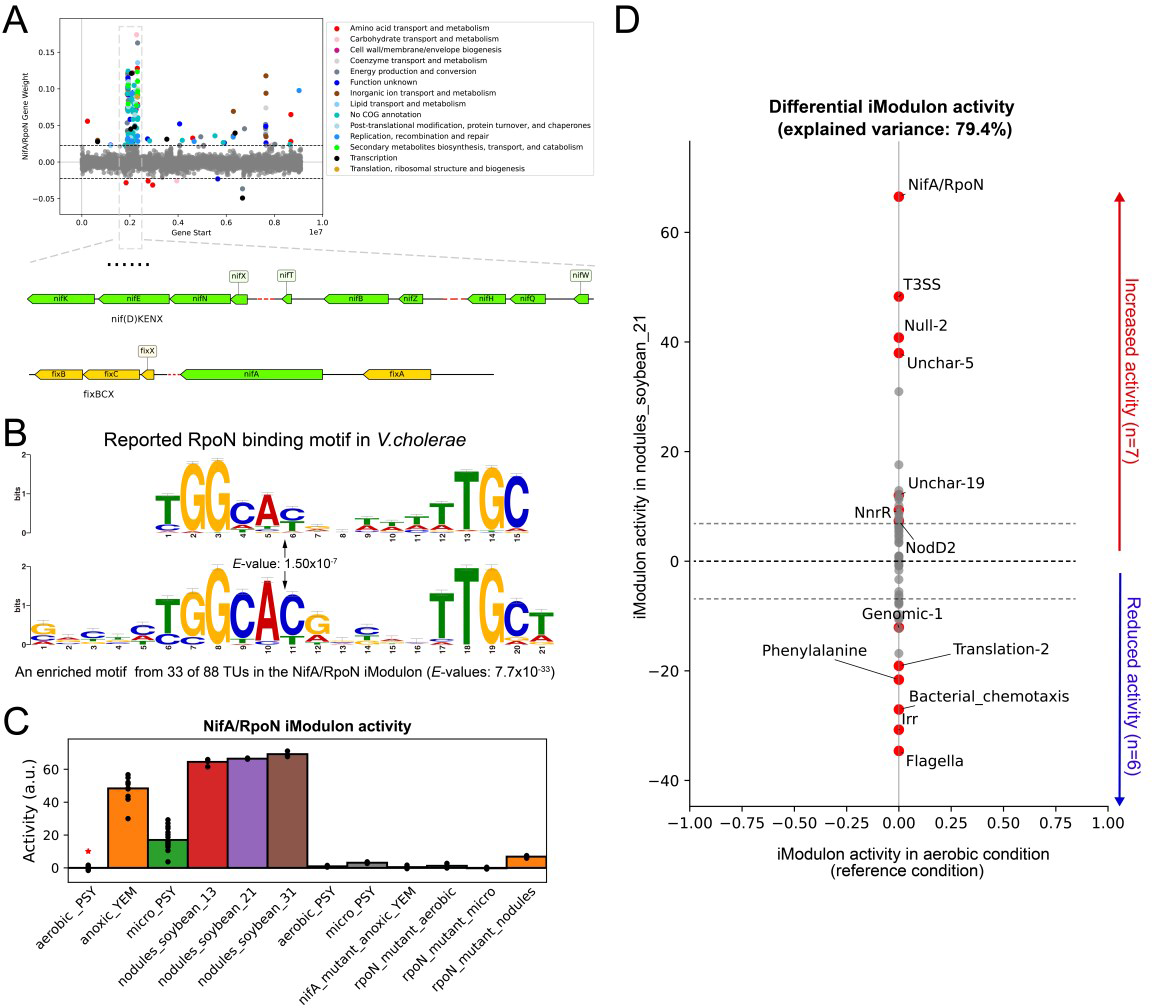
The NifA/RpoN iModulon successfully captures the nitrogen-fixing gene cluster. (A) Scatter plot of the NifA/RpoN iModulon gene weights. Genes are colored by COG categories. (B) Comparison of a motif found in the upstream regions of the NifA/RpoN iModulon genes with the previously known binding motif of RopN in *V. cholerae*. (C) NifA/RpoN iModulon activities under anoxic condition. (D) Differentially changed iModulon activities between symbiosis condition and aerobic condition (threshold = 5 and *FDR* = 0.01).

Two oxygen-responsive regulatory cascades (FixLJ-FixK_2_ and RegSR-NifA) exist in *B. diazoefficiens* USDA110. The activation status of the NifA/RpoN iModulon under different oxygen conditions is displayed in Figure 3C. It can be seen that the NifA/RpoN iModulon is activated under anaerobic conditions, and it is no longer activated when *regR*, *nifA*, or *rpoN* is knocked out. This indicates that the NifA/RpoN iModulon can effectively represent the regulatory function of the RegSR-NifA cascade.

By comparing the iModulon activity differences between free-living and symbiotic conditions, 13 significantly different iModulons were identified, including 7 significantly up-regulated and 6 significantly down-regulated iModulons (threshold = 5, *FDR* = 0.01). It can be seen that the NifA/RpoN and Flagella iModulons exhibited the largest activity changes (Figure 3D). The Flagella iModulon contains 46 genes, mainly including functional genes involved in flagella synthesis and assembly, which play a crucial role in the motility of *B. diazoefficiens* USDA110. Therefore, the decreased activity of this module suggests that under symbiotic conditions, the expression of flagella genes is suppressed and the motility is reduced, which is consistent with the function of *B. diazoefficiens* USDA110 in developing into a bacteroid.

### 3.4 Differential iModulon activation of *B. diazoefficiens* USDA110 under different environmental conditions

*B. diazoefficiens* USDA110 can provide nitrogen source for plants by forming a symbiotic system with plants to convert nitrogen into ammonia [50]. This process is very complex and involves multiple stages. First, the plant roots secrete flavonoid compounds, which attract and induce the release of nodulation factors from *B. diazoefficiens* USDA110 in the soil. Upon receiving the nodulation factor signal, the root hairs begin to curl and encase the *B. diazoefficiens* USDA110, forming an infection thread. The *B. diazoefficiens* USDA110 cells enter the plant cells along the infection thread. The plant root cortical cells then divide to form nodules, where the *B. diazoefficiens* USDA110 cells further differentiate into bacteroids for nitrogen fixation [51]. Over the past 20 years, transcriptome studies of rhizobia have been grouped according to specific stages of the symbiotic process analyzed: bacteroids vs. free-living bacteria, anaerobic vs. aerobic conditions, rhizosphere or flavonoid compounds vs. non-induced cultures, and denitrifying vs. anaerobic conditions [7].

We first compared the iModulon activities of *B. diazoefficiens* USDA110 under bacteroids and free-living aerobic conditions (threshold = 5, *FDR* = 0.05) and identified 38 significantly different iModulons. Then we investigated the activity changes of these iModulons during different stages of SNF. As shown in Figure 4, there were significant differences in the activation status of these iModulons at different stages of SNF. Under free-living aerobic conditions, when *B. diazoefficiens* USDA110 was induced by genistein for 8 hours, the Flagella iModulon responsible for flagella formation and assembly and the NodD_2_ iModulon related to nodulation factor were significantly activated, indicating that *B. diazoefficiens* USDA110 activated the motility for root recruitment and the ability to transmit nodulation signals to the plant. Under free-living microaerobic conditions, the NifA/RpoN and FixK_2_-2 iModulons were significantly activated, indicating that the anaerobic environment activated nitrogen fixation structural genes and related regulatory genes, while the Flagella iModulon was in a suppressed state, indicating that the rhizobia no longer needed the motility provided by flagella when nitrogen fixation genes were activated. Twenty-one days after nodulation, more iModulons including T3SS iModulon and T6SS iModulon were activated compared with those under free-living anaerobic conditions, indicating that at this stage, the bacteroids had the capability of extracellular secretion and transport to adapt to the substance exchange demands between the bacterial cells and the plant host. These three different environmental conditions correspond to different stages of SNF of *B. diazoefficiens* USDA110. The activity changes of these different iModulons revealed the modular activation of gene functions in response to environmental changes during SNF.

**Figure 4.**
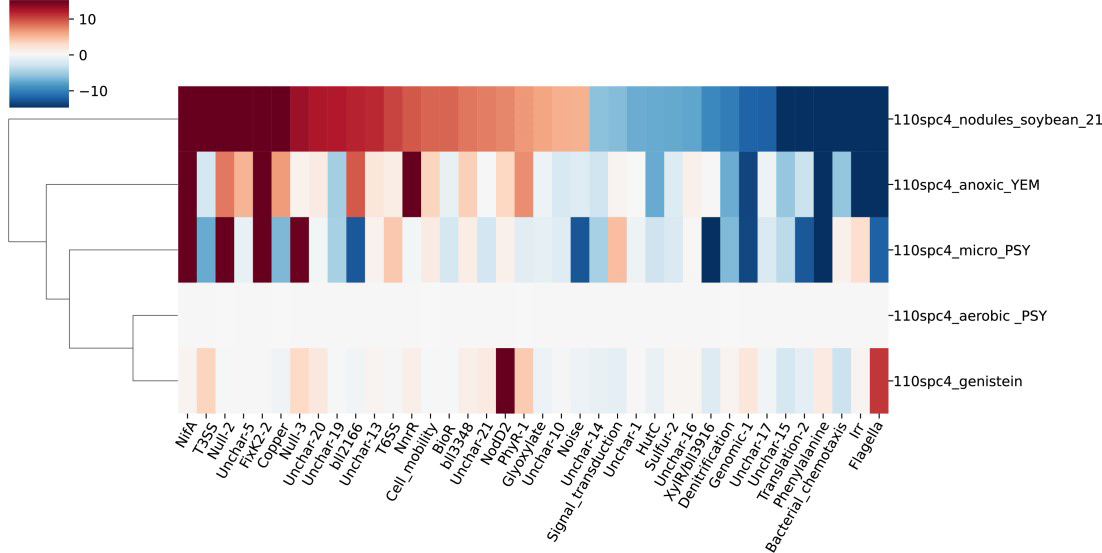
iModulon activity during different stages of symbiosis. Thirty-eight key iModulons were selected by comparing the iModulon activities of *B. diazoefficiens* USDA110 under bacteroid and free-living aerobic conditions (threshold = 5, *FDR* = 0.05). 110spc4_aerobic_PSY represents the free-living condition; 110spc4_genistein represents the state of *B. diazoefficiens* USDA110 being recruited by the plant; 110spc4_micro_PSY represents the middle stage of symbiosis; 110spc4_nodules_soybean_21 represents the late stage of symbiosis; 110spc4_anoxic_YEM represents denitrification.

### 3.5 Active-TRN based on iModulon activity

Life activities rely on the orderly regulation of gene expression, and the regulatory relationships among genes constitute the TRN. Early, to determine the regulatory relationship between genes required methods such as electrophoretic mobility shift assay, DNase I footprinting assay, and *in vitro* run-off transcription assay to verify whether TFs directly bind to the upstream promoter regions of target genes. Although these methods produced accurate results, they were time-consuming and labor-intensive, and only a small number of regulatory relationships could be captured in one experiment. Later, high-throughput technologies such as transcriptomics, proteomics, and chromatin immunoprecipitation sequencing, combined with computational analysis methods, can more easily obtain a global transcriptional regulatory landscape of a species. As is known to all, most regulatory relationships are condition-dependent, which means that the regulatory effects of TFs vary with environmental conditions [52]. However, the current research on TRN often depicts static TRNs that do not distinguish between regulatory activation states, and thus cannot describe the transcriptional regulatory relationships under different environmental conditions. To overcome this drawback, this study analyzed the regulatory effects of TFs in *B. diazoefficiens* USDA110 under four different conditions: genistein induction, microaerobiosis, bacterioid, and denitrification, based on the activity characteristics of iModulons.

As shown in the Figure 5, the active transcriptional regulatory network (active-TRN) induced by genistein contains 59 edges and 11 nodes (Figure 5A), accounting for 9.1% of the total regulatory relationships, indicating that only a small number of key pathways are activated in *B. diazoefficiens* USDA110 after receiving the genistein signal. The active-TRN under microaerobic condition contains 312 edges and 23 nodes (Figure 5B), mainly including the FixLJ-FixK_2_-FixK regulatory cascade and the NifA regulon related to low-oxygen adaptation, which is consistent with previous studies and with the related iModulon results analyzed above. The active-TRN under symbiotic conditions contains 364 edges and 30 nodes (Figure 5C), with 22 nodes appearing in both microaerobic and symbiotic conditions, and the remaining differentially expressed nodes reflect the condition-specific activation of regulatory relationships under symbiotic condition. The active-TRN under denitrification conditions contains 358 edges and 24 nodes (Figure 5D), with 21 nodes overlapping with the nodes under microaerobic condition. These results demonstrated the regulatory role of TFs in *B. diazoefficiens* USDA110 under different environmental conditions.

**Figure 5.**
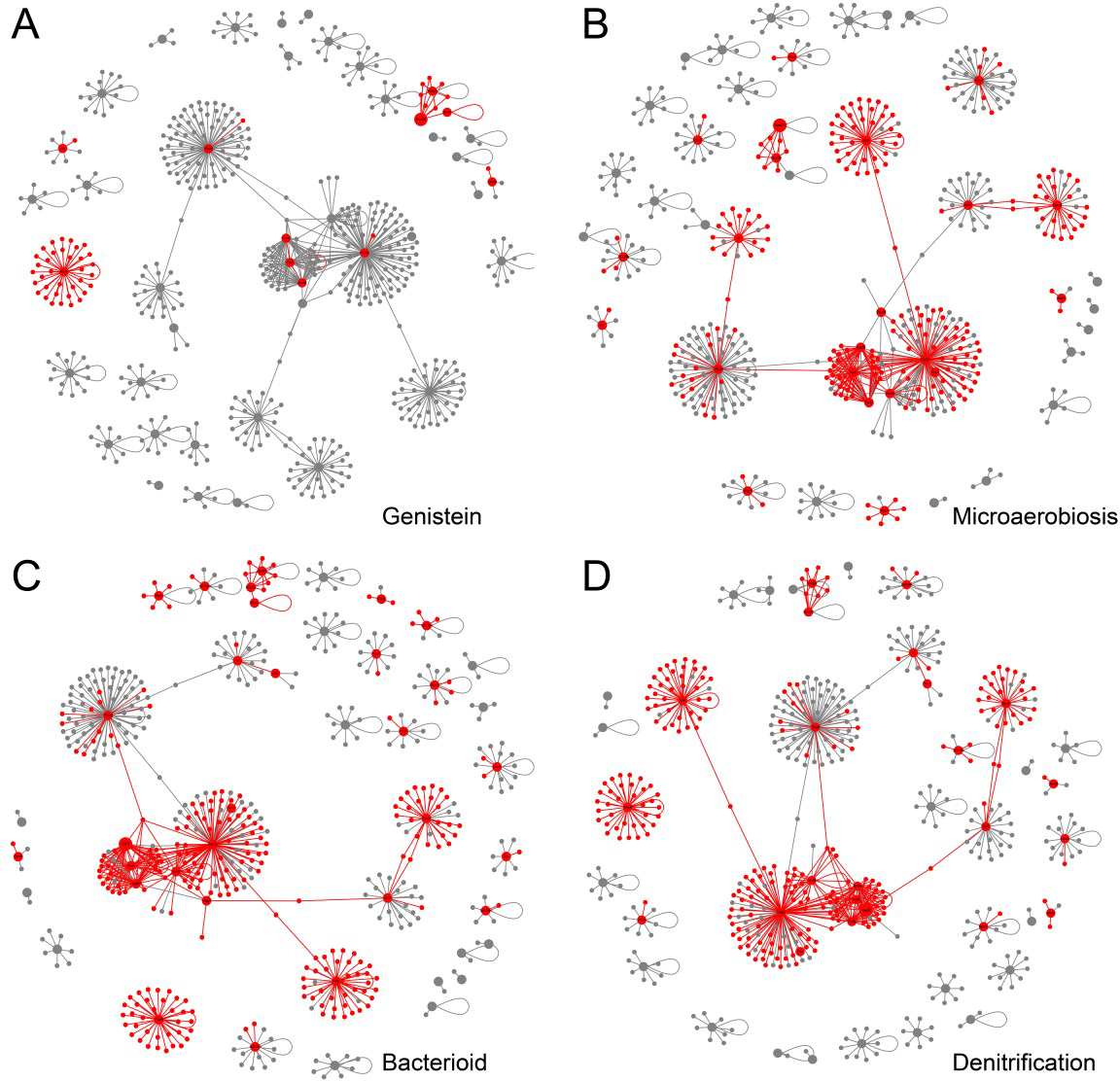
The active-TRNs based on iModulon activities. Red nodes represent active genes. (A) TRN activation after genistein induction. (B) TRN activation in microaerobiosis condition. (C) TRN activation 21 days after symbiotic nodulation. (D) TRN activation under denitrification condition.

## 4. Discussion

In this study, we used ICA to decompose 226 different transcriptomic datasets of *B. diazoefficiens* USDA110 into 64 iModulons. Many of these iModulons correspond to important TFs in *B. diazoefficiens* USDA110. Using these iModulons, we provided new insights into the relationship between the FixLJ-FixK_2_-FixK_1_ regulatory cascade and quorum sensing. We described the independence of FixLJ-FixK_2_-FixK_1_ and NifA/RpoN regulatory cascades in response to oxygen environments. We calculated the differential activation status of iModulons in *B. diazoefficiens* USDA110 under different environmental conditions, and based on this, we proposed the active-TRNs, which demonstrated the relationship between gene regulation and environmental condition.

Up to now, there are only 15 samples from 3 studies of *B. diazoefficiens* USDA110 RNA-seq data in the GEO database, which is much smaller than the amount of microarray data and is not sufficient for ICA analysis. Previous studies have found that ICA performs poorly on microarray data [53]. However, we obtained good results this time because we used data from the same microarray platform, and most of the data came from the same institution, thus avoided noise and errors introduced by differences in microarray data processing methods.

Using the ICA method, we obtained 64 iModulons from the transcriptome expression matrix of *B. diazoefficiens* USDA110, of which 13 iModulons significantly overlapped with regulons, reflecting the known regulatory relationships. Although the remaining iModulons cannot be directly associated with known regulatory relationships, most of their features can be supplemented through gene functional annotation. Among these 13 regulatory iModulons, only the HutC iModulon had a high degree of overlap with the HutC regulon and other iModulons not, possibly due to incomplete information in the TRN of *B. diazoefficiens* USDA110. The sizes of the iModulons we obtained were almost all larger than those of the corresponding regulons, indicating that an iModulon may contain multiple regulons with similar expression patterns. Through motif analysis, we discovered potential TFBSs, and the relationships between these binding sites and TFs need further verification. Although the Null-2 iModulon in our results does not contain any gene (namely, no gene has a weight above the threshold), its contribution to expression variance ranks third, which means we cannot ignore this "strange" independent signal. We found that the gene weight distribution in Null-2 iModulon was relatively average, which means that no gene significantly contributed to this independent signal. We found that the Null-2 iModulon activity significantly increased in anaerobic environment, alkaline environment, high temperature environment, and starvation environment, indicating that the Null-2 iModulon responded to extreme environments. In particular, the Null-2 iModulon activity was significantly higher in starvation than in other environmental conditions. This indicates that many genes are involved in the adaptation of the organism to extreme environments and these genes have diverse expression patterns.

By comparing the differences between FixK_2_-1 iModulon and FixK_2_-2 iModulon, we found that FixK_2_-1 iModulon is closely related to quorum sensing, while FixK_2_-2 iModulon is more representative of the FixLJ/FixK_2_/NnnR cascade. Under stress conditions, the activity of FixK_2_-1 iModulon increased, suggesting a potential connection between quorum sensing genes and the stress resistance mechanism of *B. diazoefficiens* USDA110. Studies have shown that the LuxR homologous genes in *Sinorhizobium meliloti* contribute to its adaptation to environmental challenges such as high osmotic pressure, nutrient starvation, and nodule competition, thereby increasing its chances of survival in the stressful rhizosphere environment [54]. Whether the LuxR homologous gene *blr1880* in *B. diazoefficiens* has similar functions requires further investigation. By checking the activity correlations between FixK_2_-2 iModulon and Phenylalanine iModulon, we found that the FixLJ/FixK_2_ TFs serve as a bridge connecting FixK_2_-2 iModulon and Phenylalanine iModulon. It was found that the *bll0330* gene in Phenylalanine iModulon is directly regulated by FixK_2_, and the protein product of *bll0330* gene was a two-component response regulator. This suggests that the FixLJ/FixK_2_ TFs may negatively regulate the activity of the Phenylalanine iModulon by directly regulating *bll0330*, which may involve new regulatory cascades and requires further experimental verification.

The FixLJ-FixK_2_ cascade and the RegSR-NifA cascade are relatively independent but interrelated regulatory cascades in *B. diazoefficiens* USDA110 in response to oxygen concentration [47], which are reflected in iModulon activity. Under low oxygen condition, both FixLJ-FixK_2_ and NifA/RopN iModulons show high activities. However, when *fixJ* or *fixK_2_* is absent, FixK_2_ iModulon activity decreases, but NifA/RpoN iModulon activity is not affected. Similarly, when *regR* or *nifA* or *rpoN* is absent, NifA/RpoN iModulon activity decreases, but FixLJ-FixK_2_ iModulon activity is not affected, reflecting the independence of the two regulatory cascades in response to oxygen environments. We found that the LuxR homologous gene *blr1880* seems to play a special role in regulating FixLJ-FixK_2_ iModulon activity. When *blr1880* is absent, FixLJ-FixK_2_ iModulon remains activated under aerobic conditions, while NifA/RpoN iModulon is inhibited. This indicates that *blr1880* responds to the inhibitory effect of aerobic environment on FixK_2_ iModulon activity by participating in the FixLJ-FixK_2_ cascade, but the specific mechanism still needs further verification.

We compared the activation status of iModulons under different environmental conditions and found significant differences in activities, which were consistent with the functional requirements of adaptation to the corresponding environments. In *B. diazoefficiens* USDA110, genistein induced the activation of NodD_2_ and Flagella iModulons, but did not affect other nitrogen fixation structural genes. Although nitrogen fixation-related modules were significantly activated under anaerobic conditions, additional iModulons were required for complete SNF function. Under anaerobic conditions with YEM medium, denitrification function of *B. diazoefficiens* USDA110 was induced. We found that the NifA/RpoN iModulon and FixK_2_-2 iModulon still exhibited high activities, indicating that oxygen concentration is a direct factor regulating the activities of these two iModulons. These results demonstrate the perfect coordination between gene expression and environmental adaptation in *B. diazoefficiens* USDA110, as well as the relative independence of its gene functional modules, which may find its application in synthetic biology: when the biological functions from *B. diazoefficiens* USDA110 are needed, the independence of gene functional modules benefits their implementation in other chassis organisms.

In the past decades, a series of methods have been developed to construct gene regulatory networks. The existing inference methods are based on different principles [55], and the rise of machine learning has opened up new possibilities for inferring TRNs. It should be noted that living organisms have stimulus-response coupling mechanisms to perceive and respond to changes in different environmental conditions, such as the bacterial one-component and two-component regulatory systems. These systems activate downstream regulatory cascades only after receiving specific environmental signals. However, most of the existing TRNs cannot describe the corresponding relationship between environmental condition and TF regulation. Therefore, in this study, we proposed the active-TRNs under different environmental conditions based on iModulon activity data. We assumed that when the iModulon activity significantly increased, the regulatory relationships involved in it were activated. We first identified the iModulons with significantly increased activity under different environmental conditions, and then extracted the corresponding genes and regulatory relationships. It should be noted that we did not make further screening based on changes in TF expression, because the TFs of two-component regulatory systems do not directly depend on their gene expression changes but on the proteins that sense environment signals in the system. The PyModulon software package provides an interface for inferring iModulon activity in a new dataset using a precalculated IcaData object, so with our modified scripts people can easily calculate the significantly activated iModulons in the new dataset and obtain the active-TRNs under different environmental conditions. Our results well demonstrated the TF regulatory relationships activated under different environmental conditions and extended the application of iModulon activity data.

## Supplementary materials

**Figure S1.**
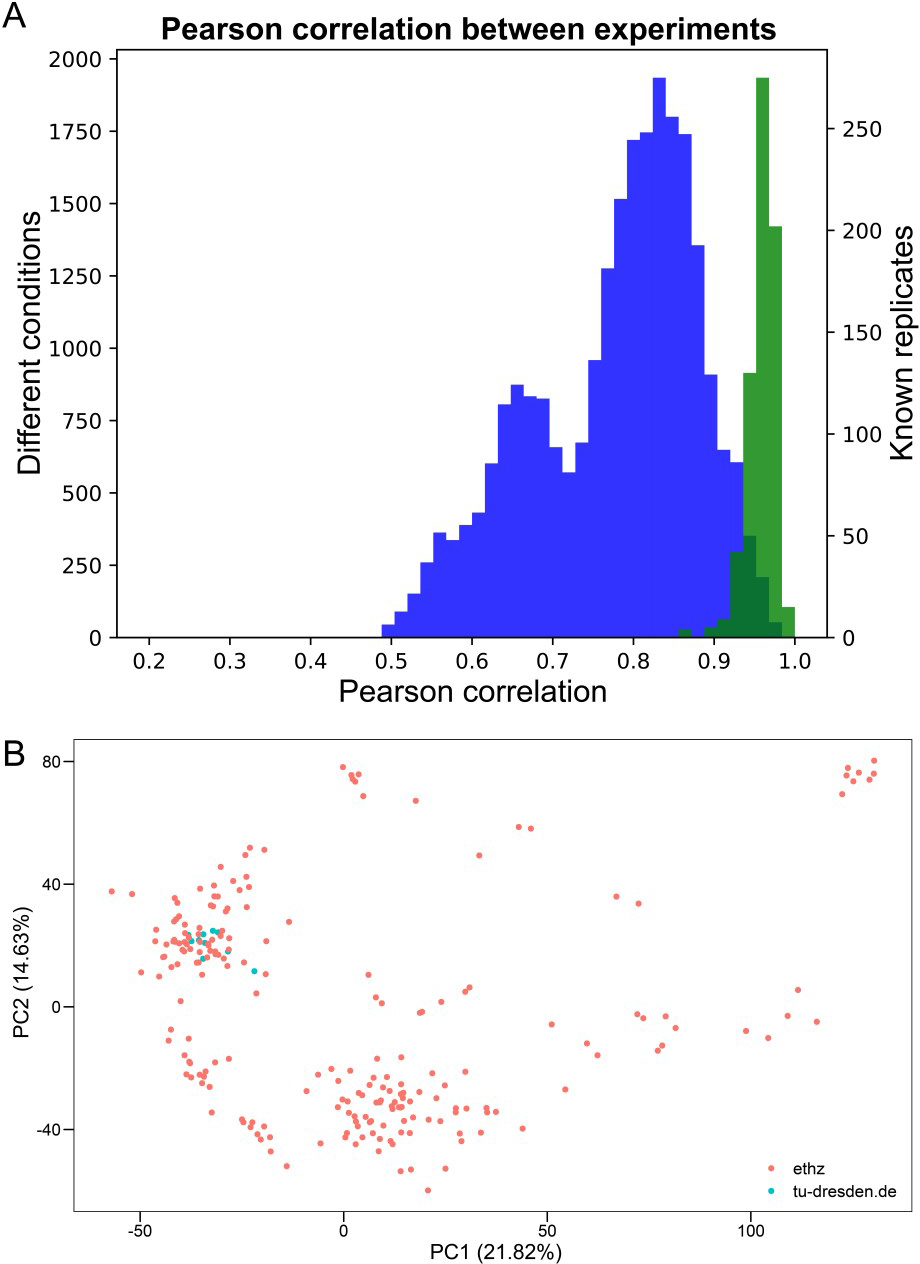
Summary of data. (A) Histogram of the Pearson correlation coefficient (PCC) between gene expression data, after centering to reference condition. (B) Loadings of the first two principal components (PC) of the compendium, colored by the researchers’ institution. Red represents ethz (ETH Zurich) and blue represents tu-dresden (Dresden University of Technology). There were 216 samples from ethz or with ethz involved, and 10 samples from tu-dresden. The high coincidence of red and blue indicates high self-consistency.

**Figure S2.**
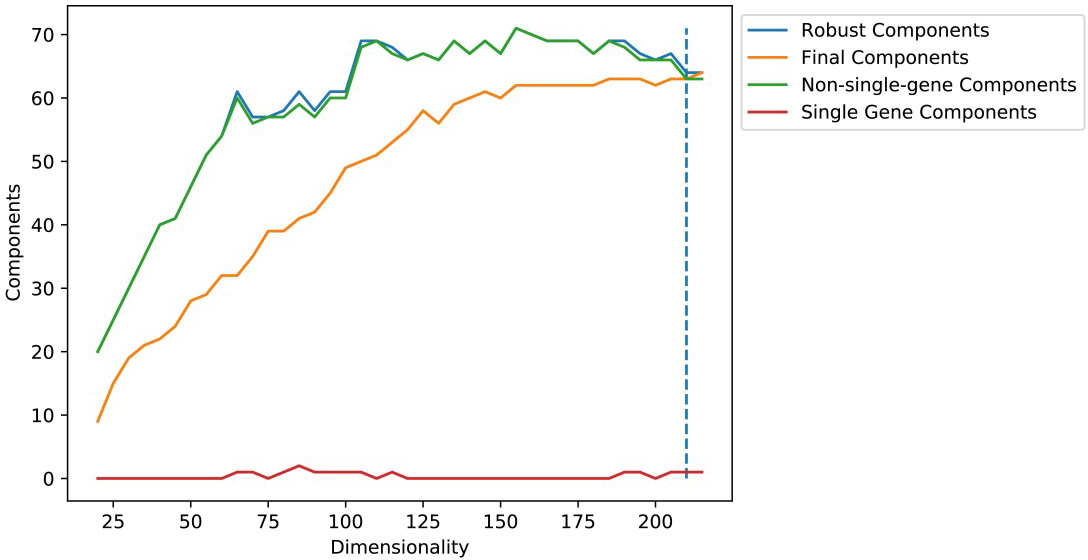
Dimension analysis. ICA was used in gene expression datasets and some independent components were obtained. As mentioned in **Materials and methods**, we performed clustering of the gene expression profile multiple times for dimensions between 20 and 215 with a step size of 5. All components computed from a multi-start ICA decomposition were counted as “robust components”. A component was classified as “single gene” if the highest gene weight was more than twice the next highest; the number of non-single gene components was determined by subtracting the number of single gene components from the number of robust components [31].

**Figure S3.**
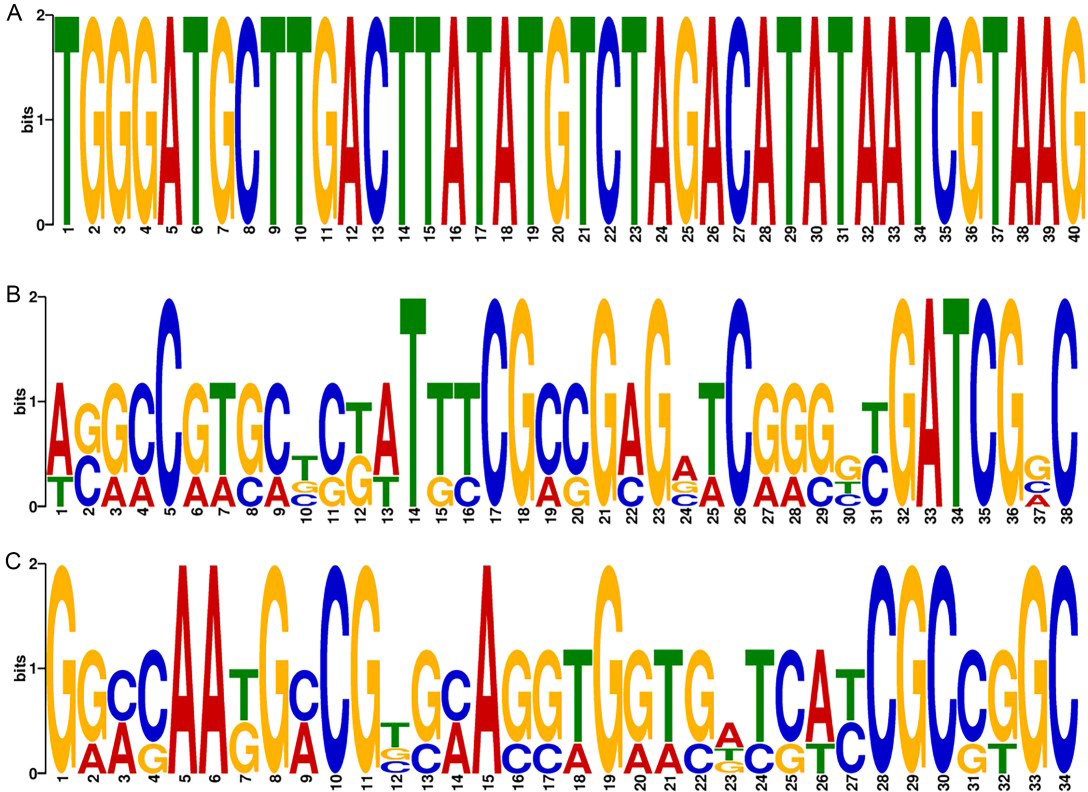
The motifs identified in the upstream promoter regions of HutC iModulon genes. (A) Motif-1. (B) Motif-2. (C) Motif-3.

**Figure S4.**
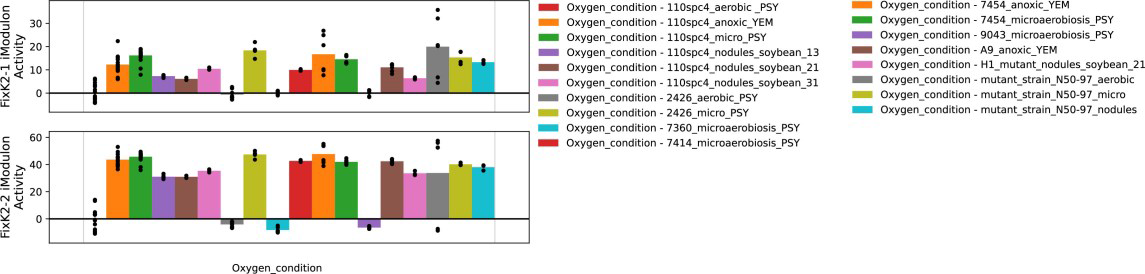
The activities of FixK_2_-1 iModulon and FixK_2_-2 iModulon in response to oxygen conditions. 7360_microaerobiosis_PSY represents the 110spc4 *fixJ* mutant and 9043_microaerobiosis_PSY represents the 110spc4 *fixK_2_*mutant.

**Figure S5.**
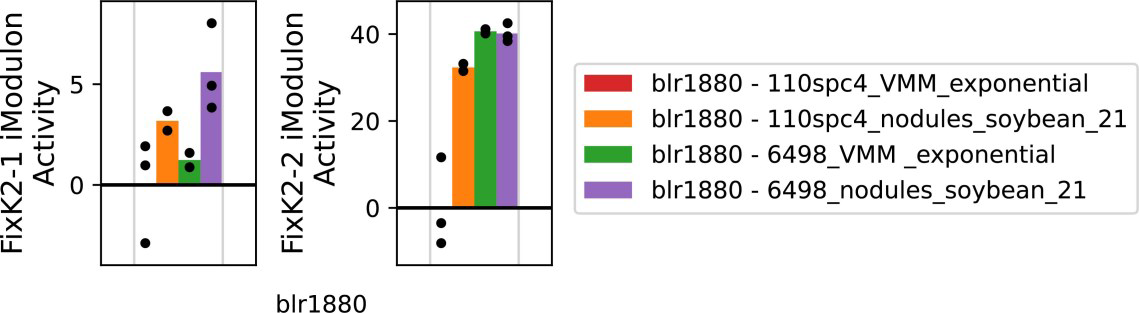
FixK_2_-1 iModulon and FixK_2_-2 iModulon activities in the *blr1880* project. 6498_VMM_exponential represents the *blr1880* mutant growing in VMM minimal medium and 6498_nodules_soybean_21 represents the *blr1880* mutant growing with soybean.

